# Splicing factor BUD31 promotes ovarian cancer progression through sustaining the expression of anti-apoptotic *BCL2L12*

**DOI:** 10.1101/2022.01.19.476862

**Authors:** Zixiang Wang, Shourong Wang, Junchao Qin, Xiyu Zhang, Gang Lu, Hongbin Liu, Haiyang Guo, Ligang Wu, Changshun Shao, Beihua Kong, Zhaojian Liu

## Abstract

Dysregulated expression of splicing factors has important roles in cancer development and progression. However, it remains a challenge to identify the cancer-specific splicing variants. Here we demonstrated that spliceosome component BUD31 is increased in ovarian cancer, and its higher expression predicts worse prognosis. We characterized the BUD31 binding motif and found that BUD31 preferentially binds exon-intron regions near splicing sites by CLIP-seq. Further analysis revealed that BUD31 inhibition results in extensive exon skipping and decreased abundance of long CDS isoforms. In particular, we identified *BCL2L12*, an anti-apoptotic BCL2 family member, as a functional splicing target of BUD31. BUD31 stimulates the inclusion of exon 3 to generate full-length *BCL2L12* and promotes ovarian cancer progression. Knockdown of BUD31 or splice-switching antisense oligonucleotide treatment promotes exon 3 skipping and results in a truncated isoform of BCL2L12 that undergoes nonsense-mediated mRNA decay, and the cells subsequently undergo apoptosis. Our findings reveal BUD31-regulated exon inclusion as a critical factor in ovarian cancer cell survival and progression.

## INTRODUCTION

Serous ovarian carcinoma (SOC) is the most prevalent subtype and accounts for ∼85% of ovarian cancer (Zhang et al., 2016). High-grade serous ovarian carcinoma (HGSOC) is aggressive, with only about a 30% five-year survival rate (Kim et al., 2018). HGSOC is traditional thought to development from ovarian surface epithelium (Ducie et al., 2017). Since the late 1990s, accumulating evidence indicates that HGSOC originates from fallopian tube secretory epithelial cells (DeWeerdt, 2021; Przybycin et al., 2010). However, recent studies demonstrate that both fallopian tube and ovarian surface epithelium are origin of HGSOC (Lohmussaar et al., 2020; Zhang et al., 2019). Identification of the cell of origin of HGSOC still remains a challenge. PARP inhibitors have shown promising clinical results for the treatment of ovarian cancer (Chen and Du, 2018). Despite advances in understanding molecular aberrations and the target therapy of ovarian cancer, relapse and chemotherapy resistance is a major clinical problem.

Alternative splicing (AS) of precursor mRNA is an important step to increase the diversity of gene expression, and AS has been estimated to occur in about 95% of human multi-exon genes (Scotti and Swanson, 2016). While AS is essential for normal development, dysregulation of the splicing process is implicated in various diseases, including cancer (Leclair et al., 2020), and malignant tumors have up to 30% more AS events than normal tissues (Kahles et al., 2018). AS is regulated by trans-splicing factors that specifically bind to cis-elements in pre-mRNAs (Matera and Wang, 2014).

Mutations or altered expression of splicing factors can lead to splicing reprogramming, which contributes to tumor initiation and progression. For example, SF3B1 mutations induce mis-splicing of *BRD9*, leading to its degradation and the promotion of melanomagenesis (Inoue et al., 2019). U2AF1 mutations cause abnormal recognition of the 3’ splice site of pre-mRNA, resulting in increased DNA damage in cancers (Wang and Aifantis, 2020). SRSF1 is overexpressed in various cancers and exerts oncogenic roles by regulating the AS of genes, including MYO1B (Du et al., 2021; Luo et al., 2017; Zhou et al., 2019). The splicing factor ESRP1 regulates *CD44* splice switching during the epithelial-mesenchymal transition (Brown et al., 2011), and SF3B2 drives prostate cancer progression through AS of androgen receptor (AR) to increase AR-V7 expression (Kawamura et al., 2019). Conversely, RBM4 is downregulated in cancer tissues, and this suppresses tumor progression by modulating *Bcl-x* splicing (Wang et al., 2014). Dysregulation of AS provides a novel therapeutic strategy for cancer treatment. For example, H3B-8800 is an SF3B1 inhibitor, which is currently in a phase I trial for the treatment of various cancers (Steensma et al., 2021), and antisense oligonucleotides (ASOs) for modulating pre-mRNA splicing are currently FDA-approved for the treatment of spinal muscular atrophy (El Marabti and Abdel-Wahab, 2021). ASOs modulate *Bcl-x* splicing to produce the pro-apoptotic isoform and reduce the tumor load in mice (Bauman et al., 2010), suggesting a promising approach for cancer therapy. Multiple AS markers have been identified in ovarian cancer (Klinck et al., 2008), including aberrant expression of splicing factors such as SRSF3 and SFPQ (He and Zhang, 2015; Pellarin et al., 2020). Hyperactivation of MYC can lead to global upregulation of pre-mRNA levels and to aberrant splicing patterns in ovarian cancer (Anczukow and Krainer, 2015). However, knowledge of the role of splicing factors in the generation of ovarian cancer-related splicing variations is still limited.

BUD31 is a spliceosomal component in yeast (Masciadri et al., 2004), which is required for spliceosome assembly and catalytic activity (Hsu et al., 2015). BUD31 is identified as a MYC-synthetic lethal gene in human mammary epithelial cells (Hsu *et al*., 2015), implying its potential role in cancer. Nonetheless, the alternative splicing regulation and clinical significance of BUD31 in cancer remain poorly understood. In this work, we report that overexpression of BUD31 predicts poor prognosis in ovarian cancer patients. Furthermore, we have used RNA-seq and CLIP-seq analysis to show BUD31-regulated AS and identify the binding motif and preferred genome-wide binding pattern of BUD31. More importantly, we verified that BUD31 drives an oncogenic splicing switch of *BCL2L12*, which in turn promotes ovarian cancer progression. Our study indicates that BUD31 is a critical oncogenic splicing factor that might act as a potential therapeutic target in ovarian cancer.

## RESULTS

### Elevated BUD31 expression is associated with poor prognosis in ovarian cancer

To identify survival-related splicing factors in SOC, we first analyzed the expression of 134 known splicing factors (Wang et al., 2021) in SOC tissues (n = 374) compared to normal tissues (n = 180) from the TCGA and GTEx databases. We identified 20 upregulated and 17 downregulated splicing factors in SOCs (Figure 1A). We next assessed the prognostic values of 37 dysregulated splicing factors in patients with SOC and found 10 of 37 splicing factors to be significantly related to both progression-free survival and overall survival (Figures 1B and 1C). Among these, BUD31 caught our attention because overexpression of BUD31 was related to poor prognosis in ovarian cancer patients. We further examined the expression level of BUD31 in the TCGA and Clinical Proteomic Tumor Analysis Consortium (CPTAC) data. BUD31 was commonly upregulated in SOC compared with normal samples at both the RNA and protein level (Figures 1D and 1E). Importantly, BUD31 protein level was significantly increased in advanced ovarian cancer compared to early-stage patients (Figure 1E). Pan-cancer analysis revealed that BUD31 is overexpressed in various cancer types. To evaluate the clinical significance of BUD31 in SOC, we performed immunohistochemistry using a tissue microarray containing 149 ovarian cancer tissue samples and 73 fallopian tube tissues (FTs). Immunohistochemical (IHC) staining revealed the significant overexpression of BUD31 in SOCs (Figures 1F and 1G). We further assessed the prognostic value of BUD31 in SOCs in this cohort and found that a high level of BUD31 was significantly correlated with poor overall survival and progression-free survival in patients with SOC (Figures 1H and 1I), verified by the Kaplan–Meier Plotter database (Figures 1J and 1K). Moreover, a high level of BUD31 was positively associated with being younger than 60 years old. Together, these results demonstrate that elevated expression of BUD31 is associated with worse prognosis in ovarian cancer patients.

**Figure 1.**
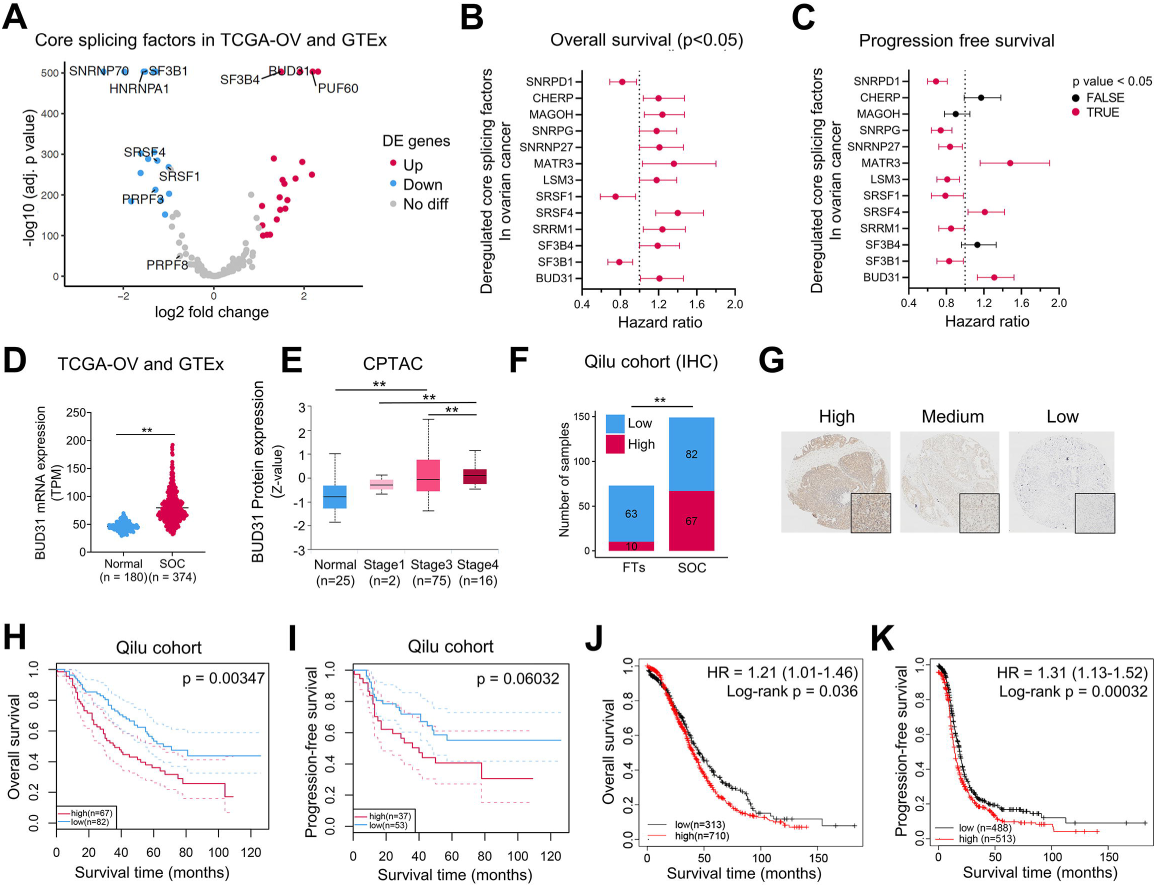
High BUD31 expression is associated with poor prognosis in ovarian cancer. (A) Volcano plot of differentially expressed core splicing factors (n = 134) between the TCGA-OV cohort (n = 374) and normal tissue in GTEx datasets (n = 180). |log2FC| > 1 and an adjusted p-value < 0.05 were considered significant. (B and C) Forrest plot of the hazard ratio for the association between 13 dysregulated splicing factors and overall survival (B) and progression-free survival (C) in patients with SOC from the Kaplan–Meier Plotter database. The cohort of patients with SOC was split by median expression values through the autoselect cutoff. Prognostic splicing factors (red) with a p < 0.05 were considered statistically significant. (D) *BUD31* mRNA expression was analyzed in the TCGA-OV cohort (n = 374) and normal tissue in GTEx datasets (n = 180). P-value was calculated from a two-tailed unpaired Student’s t-test. (E) BUD31 protein level was analyzed in SOCs (n = 93) and FTs (n = 25) from the CPTAC dataset. SOC samples were classified into Stage1 (n = 2), Stage3 (n = 75), and Stage4 (n = 16) according to individual cancer stage. Z-values represent standard deviations from the median across samples for the given cancer type or stage. P-values were calculated from a two-tailed unpaired Student’s t-test. (F) Statistical analysis of BUD31 expression from IHC staining of the tissue microarray containing 149 samples of SOCs and 73 samples of FTs collected from Qilu Hospital, Shandong University. Qilu cohort of patients with ovarian cancer was dichotomized into high (Score ≥ 7) and low (Score < 7) expression groups according to their IHC staining score as described (Wang *et al*., 2021). P-value was calculated from the Chi-square test. (G) Representative images of IHC staining with high, medium, and low BUD31 expression in our tissue microarray. (H-I) Kaplan–Meier analysis of the correlation between BUD31 expression and overall survival (H) and progression-free survival (I) of ovarian cancer patients based on data from our tissue microarray. The 95% confidence interval of high (red) and low (blue) expression groups defined by IHC staining score were shown as dotted lines. All patients with overall survival or progression-free survival information were included. (J-K) Kaplan–Meier analysis of the correlation between BUD31 expression and overall survival (J) and progression-free survival (K) of ovarian cancer patients based on the Kaplan–Meier Plotter cohort. The cohort of patients with serous ovarian cancer was split by median expression values through autoselect cutoff, and the p-value was obtained using log-rank test. Data are presented as means ± S.D. *p < 0.05, **p < 0.01.

### Knockdown of BUD31 induces spontaneous apoptosis in ovarian cancer cells

To investigate the functional role of BUD31 in ovarian cancer, we established stable cell lines with BUD31 overexpression (A2780 and OVCAR3) or dox-inducible BUD31 knockdown (HEYA8) relative to the basal expression level of BUD31. We then performed RNA-seq on BUD31 knockdown and control HEYA8 cells and identified 1,243 downregulated and 943 upregulated genes. Among the downregulated genes upon BUD31 knockdown, 31.86% were oncogenic genes highly expressed in SOCs. Gene Ontology (GO) analysis revealed that BUD31 target genes were enriched in biological processes, including apoptosis, cell division, and microtubule cytoskeleton organization (Figures 2A and 2B). Consistent with this, the gene set enrichment analysis (GSEA) demonstrated that BUD31 knockdown regulates the apoptosis signaling pathway (Figure 2C). We next measured apoptosis in HEYA8 cells upon BUD31 knockdown using Annexin V/7-AAD staining and flow cytometry. As expected, knockdown of BUD31 induced significant spontaneous apoptosis in HEYA8 and OV90 cells (Figures 2D, left panel). The expression of apoptosis-related proteins was next measured by western blot, and inactivation of BUD31 resulted in an increased Bax/Bcl-2 ratio along with increased cleaved caspase-3 and PARP1 (Figure 2E, left panel). Additionally, BUD31 knockdown using siRNAs resulted in depolymerized microtubules and abnormal cell morphology in HEYA8 cells. Consistent with this, overexpression of BUD31 suppressed H_2_O_2_-induced ovarian cancer cell apoptosis (Figures 2D and 2E, right panel). These results indicate that high levels of BUD31 exert an anti-apoptosis effect in ovarian cancer.

**Figure 2.**
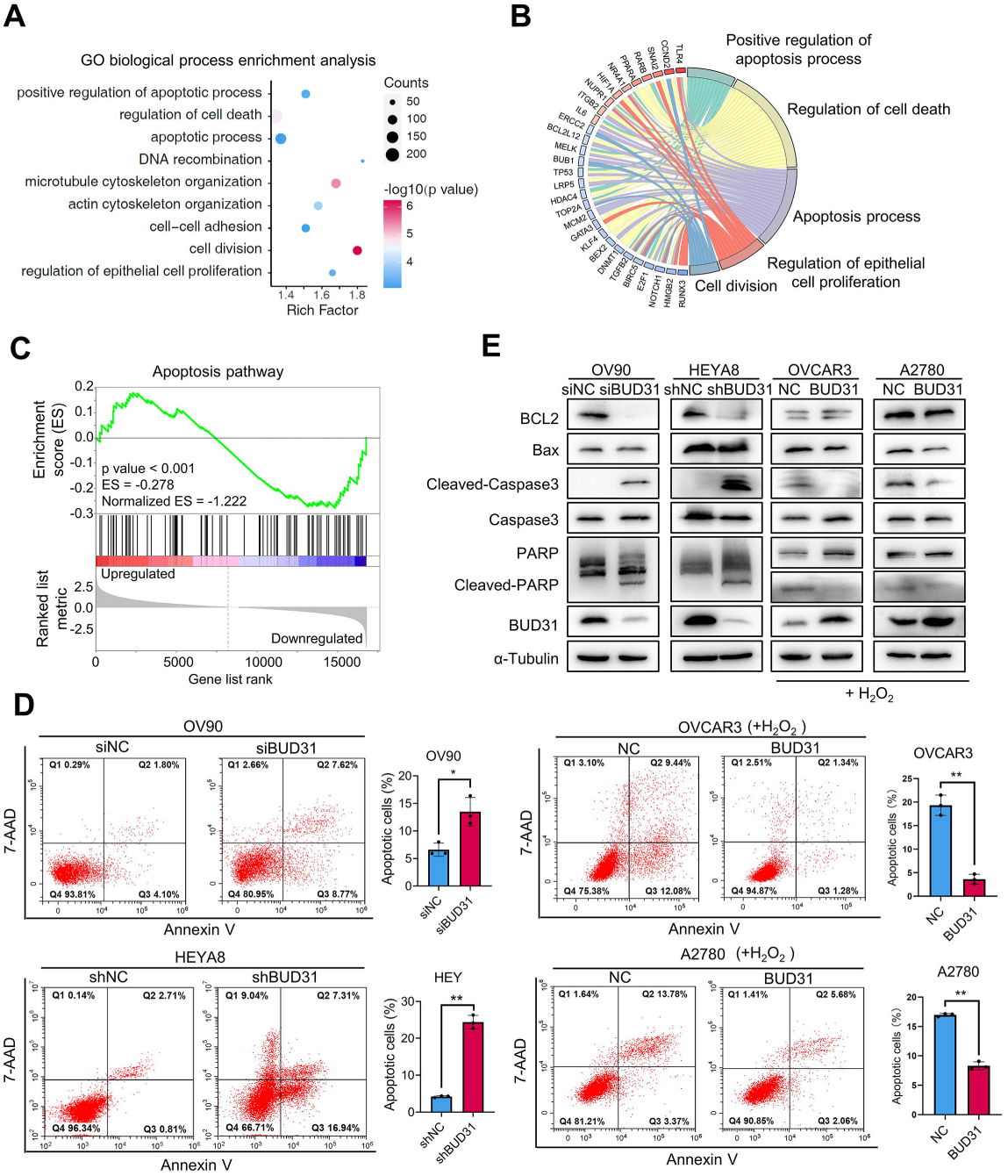
Knockdown of BUD31 induces spontaneous apoptosis in ovarian cancer cells. (A-B) GO biological process enrichment (A) and Circos plot (B) analysis were conducted on DEGs in the RNA-seq data from HEYA8 cells after BUD31 knockdown. (C) GSEA analysis was performed with the gene expression profile after BUD31 knockdown (Normalized ES = – 1.222, p < 0.001). (D) Apoptotic cells were detected by flow cytometry after staining with Annexin V/7-AAD in ovarian cancer cells after BUD31 knockdown (HEYA8 and OV90) or overexpression (OVCAR3 and A2780). Apoptotic cells percentage included early and late apoptotic cells. Cells overexpressing BUD31 were treated with H_2_O_2_ with a final concentration of 400 μM for 4 h before apoptosis detection. (E) Apoptotic markers were measured by western blot. BUD31 was knocked down in HEYA8 and OV90 cells and was overexpressed A2780 and OVCAR3 cells in the presence of H_2_O_2_. The p-value was obtained by two-tailed unpaired Student’s t-test (D), and data are presented as means ± S.D. *p < 0.05, **p < 0.01

### BUD31 promotes proliferation and xenograft tumor growth in ovarian cancer

To further explore the function of BUD31 in ovarian cancer, we performed an EdU (5- ethynyl-2′-deoxyuridine) assay and found that BUD31 overexpression in A2780 and OVCAR3 cells significantly increased the number of EdU-positive cells. In contrast, knockdown of BUD31 in OV90, OVBWZX (primary ovarian cancer cells derived from the ascites of SOC patients, verified by PAX8 and p53), and HEYA8 cells reduced the number of EdU-positive cells (Figures 3A). Additionally, growth curve and clonogenic assays showed that BUD31 overexpression significantly enhanced the proliferation of ovarian cancer cells, while silencing BUD31 had the opposite effect (Figures 3B). Moreover, mouse xenograft experiments were conducted to assess the functional role of BUD31 in ovarian cancer tumorigenesis and progression. Luciferase-expressing HEYA8 cells with dox-inducible BUD31 knockdown and corresponding control cells were intraperitoneally injected into nude mice (n = 6), and luciferase was used as a tracer for in vivo imaging analysis. BUD31 knockdown in HEYA8 cells significantly reduced the number and size of tumor nodes in the abdominal cavity (Figures 3C). We also found that BUD31 knockdown reduced the Ki-67 index (Figures 3D). Furthermore, BUD31 knockdown enhanced apoptosis in xenograft tumors of HEYA8 cells as detected by TUNEL assay (Figure 3E). Therefore, these results suggest that BUD31 exhibits oncogenic potential in ovarian cancer.

**Figure 3.**
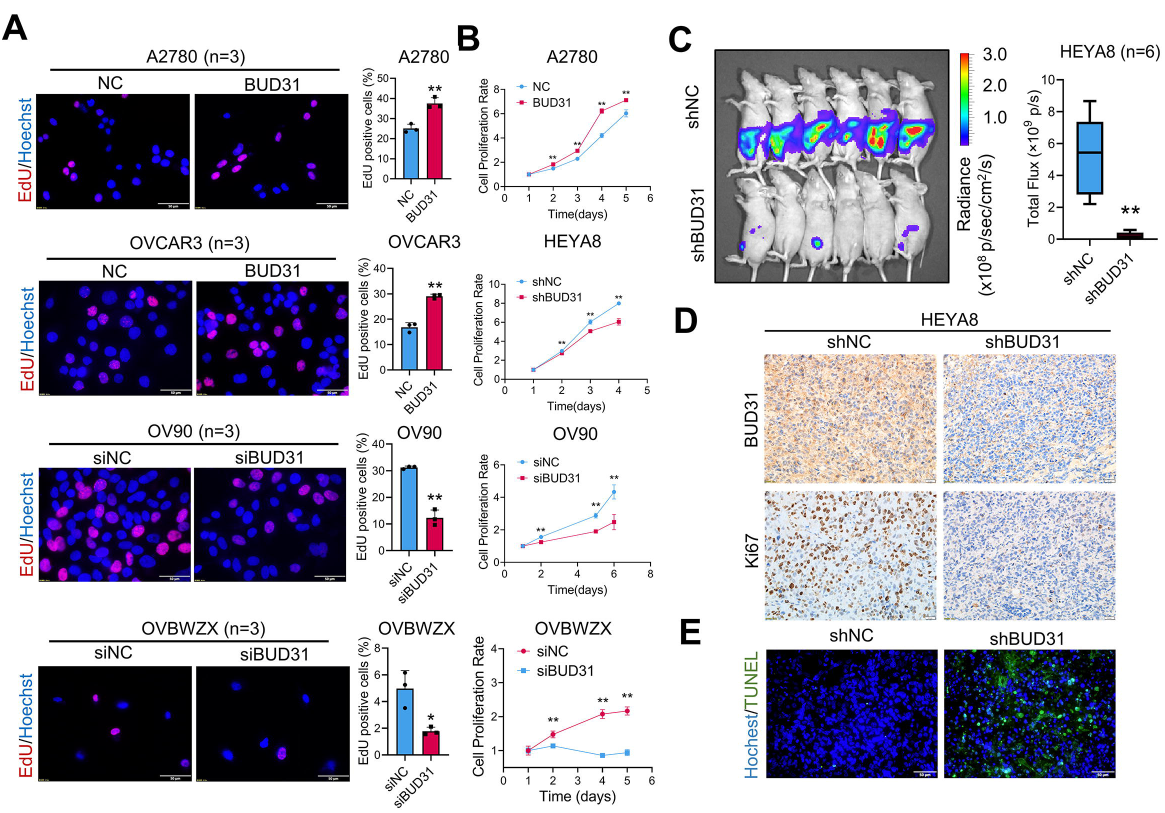
BUD31 promotes proliferation and xenograft tumor growth in ovarian cancer. (A-B) The EdU assay (A) and cell proliferation assay (B) were performed in ovarian cancer cells with BUD31 overexpression (A2780, OVCAR3) or knockdown (HEYA8, OV90, OVBWZX) compared to corresponding controls (n = 3 for the EdU and n = 5 for the cell proliferation assay). Cell proliferation was measured using the MTT cell proliferation assay. Absorbance at 570nm at each time point was compared to the initial state (time = 1 day). (C) Luciferase signals of intraperitoneal injected nude mice and photon flux quantification. Nude mice were injected with luciferase-expressing HEYA8 cells with a dox-inducible BUD31 knockdown system (n = 6 per group). Administration of doxycycline (1.2 g/L) started one week after the cell implantation. (D) IHC staining of BUD31 and Ki67 expression in xenograft tumors of HEYA8 cells with BUD31 knockdown compared to corresponding controls. (E) TUNEL assay to quantify the apoptotic cells in xenograft tumors with BUD31 knockdown compared to corresponding controls. The p-value was obtained by Student’s t-test (A, B, C), and the results are presented as the mean ± S.D. *p < 0.05, **p < 0.01.

### Identification of the genome-wide landscape of BUD31 binding sites on RNA

To unveil the role of BUD31 in AS, we first identified proteins that are associated with BUD31 by immunoprecipitation with BUD31 antibody coupled to mass spectrometry (IP-MS) in HEYA8 cells. GO enrichment analysis showed that the mRNA splicing via the spliceosome and regulation of RNA splicing pathways were significantly enriched among proteins interacting with BUD31 (Figure 4A). Intriguingly, we found that 46 annotated spliceosome proteins were associated with BUD31. Among these, BUD31 immunoprecipitated predominantly with U2 snRNPs and hnRNP proteins (Figure 4B).. Interestingly, the immunofluorescence assay showed that BUD31 was colocalized with SC35, which is a marker of the nuclear speckle (a type of nuclear body involved in splicing factor storage) (Figure 4C). The interactions between BUD31 and multiple spliceosome components suggest an essential role for BUD31 in the regulation of AS.

**Figure 4.**
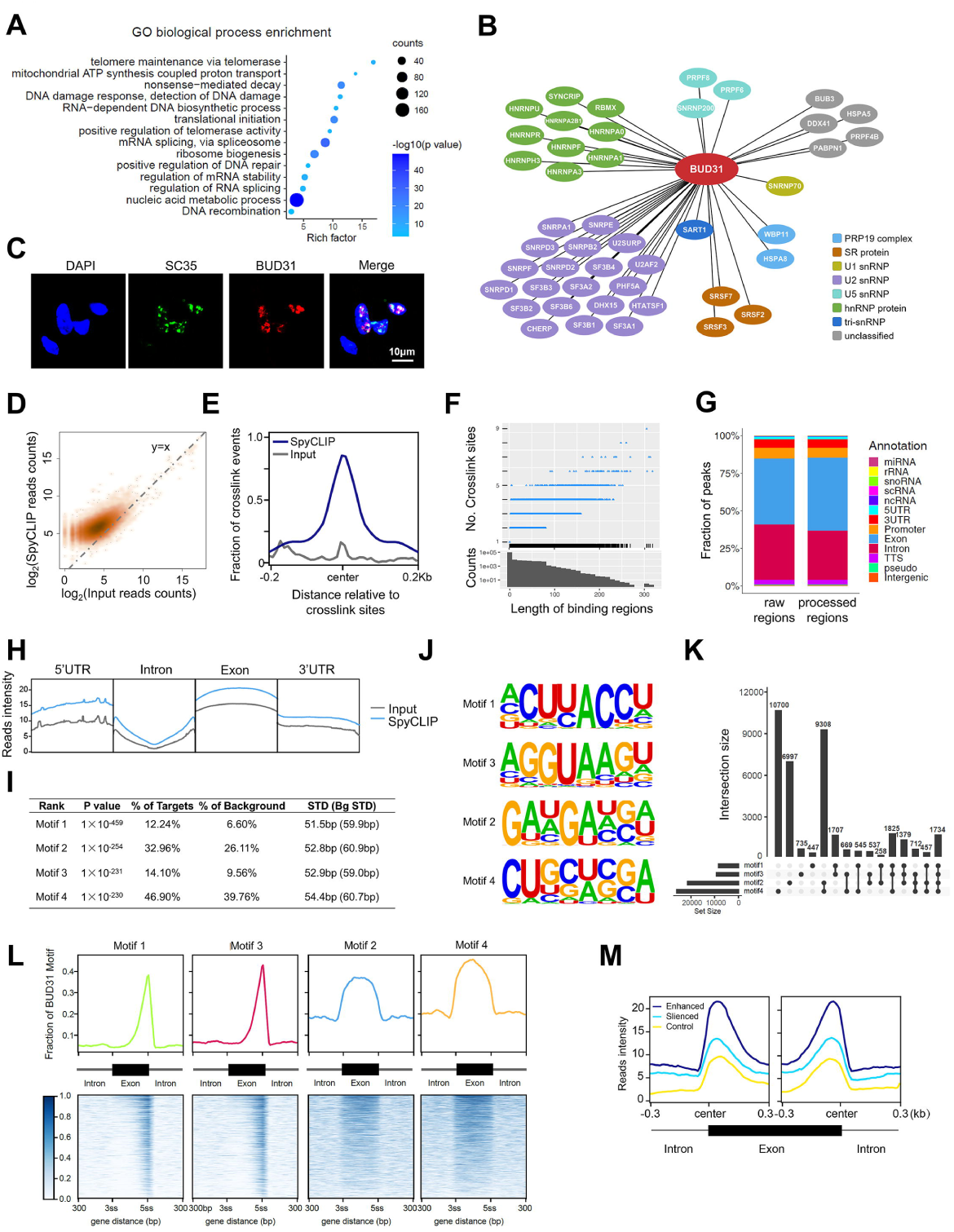
Identification of the genome-wide BUD31 binding sites on RNA. (A) GO biological process enrichment analysis of BUD31-interacting proteins in HEYA8 cells using immunoprecipitation coupled to mass spectrometry analysis with the BUD31 antibody. (B) The correlation network between BUD31 and splicing factors was constructed in Cytoscape. Proteins belonging to the spliceosome were classified into PRP19 complex, SR protein, U1 snRNP, U2 snRNP, U5 snRNP, hnRNP protein, tri-snRNP, and others. (C) Immunofluorescence assays showed the colocalization of BUD31 (red) with splicing factor SC35 (green) in punctate nuclear speckles. (D) High-density scatter plot of the SpyCLIP and input reads counts aligned to the BUD31 binding regions. (E) Position of the SpyCLIP crosslinking regions relative to the crosslinking sites identified by PURECLIP. The SpyCLIP (deep blue) and input (gray) signals are shown around the crosslink sites. (F) Length distribution of the BUD31 binding regions and the crosslinking sites included in the corresponding binding regions. Crosslinking sites whose distance from adjacent crosslinking sites was shorter than 80 nucleotides were combined into binding regions. (G) Distribution of SpyCLIP crosslinking regions annotated by HOMER on genome elements. Processed regions were longer than 3 nucleotides. (H) SpyCLIP read distribution compared with input on the genome elements, including 5’ UTR, intron, exon, and 3’ UTR. (I) De novo motif analysis of BUD31 SpyCLIP clusters and statistical results of the top-four BUD31 binding motifs. BUD31 binding motifs were ranked by p-value, and targets percentage, background percentage, and standard deviation are listed. (J) Enriched sequence elements of the top-four BUD31 binding motifs. (K) Upset plot of the distribution of motifs 1–4 in the BUD31 binding regions. (L) BUD31 binding motif distribution in the exon region and the 300 bp flanking the 3ss or 5ss intron-exon junction site. The exon region was scaled such that the length was equal to 300 bp to normalize different exon lengths. (M) SpyCLIP reads intensity distributed around the alternative exons. Enhanced exons (deep blue) were included, and silenced exons (blue) were excluded after silencing BUD31. Randomly chosen exons were used as controls (yellow).

To generate genome-wide maps of BUD31 protein-RNA interactions, we performed SpyTag-based CLIP (SpyCLIP) for BUD31 in HEYA8 cells as previously described (Zhao et al., 2019). SpyCLIP is a covalent link-based CLIP method with high efficiency and accuracy. SpyTag-SpyCatcher system could withstand harsh washing for removing non-specific interactions (Zhang et al., 2020). We got CLIP-seq data with high accuracy and reproductivity, which was suitable for further analysis. A total of 9,983,683 reads in the input group and 30,970,512 reads in the SpyCLIP group were obtained, and 99.02% of the SpyCLIP reads were mapped to an annotated human genome (hg38). Further cluster analysis revealed that BUD31 binds to 8,780 annotated human genes. Protein-RNA crosslink sites were identified by the PURECLIP method, which explicitly incorporates CLIP truncation patterns and non-specific sequence biases (Krakau et al., 2017). The identified regions shared more SpyCLIP reads than the control group, and SpyCLIP exhibited strong enrichment at the crosslink sites (Figures 4D, 4E). Most BUD31-binding regions were less than 280 nucleotides in length and contained less than six crosslink sites (Figure 4F), and more than 87.60% of the BUD31-RNA crosslink regions mapped to exons and introns (Figures 4G and 4H). To better characterize the interaction of BUD31 with RNA, we used the HOMER algorithm (Heinz et al., 2010) to identify the BUD31-recognizing RNA motif and found that the most abundant element was the ACUUACCU 8-mer (Figures 4I, 4J, and 4K). Strikingly, 2 of the 4 top-scoring motifs (motif 1 and motif 3) were located near the 5ss intron-exon junction and were reverse complemented. The other two top-scoring motifs located in exon regions (Figures 4L). We further conducted correlation analysis between BUD31 binding regions and the regulated alternative exons based on SpyCLIP and RNA-Seq data. Intriguingly, the BUD31 binding sites were highly enriched in exon-intron regions near both the 3’ and 5’ splicing sites (Figures 4M). To sum up, these results suggest that BUD31 exerts its function in AS through direct interactions with the pre-mRNA substrate.

### Global identification of AS events regulated by BUD31

To investigate BUD31-regulated AS events, AS analysis was performed based on the RNA-seq data in BUD31 knockdown and control HEYA8 cells. 3,932 AS events with IncLevelDifference (the “Percent Spliced In” (PSI) change) over 10% were identified. These included retained introns, skipped exons, alternative 5’ splice sites, alternative 3’ splice sites, and mutually exclusive exons. The predominant AS event upon BUD31 knockdown was skipped exons (68.0%) (Figures 5A). Further global AS analysis revealed that skipped exons and retained introns led to significant isoform switches and part of genes expression changes (Figures 5B, 5C). Consistent with this, the global coding sequence (CDS) length was calculated, showing that knockdown of BUD31 significantly decreased the abundance of long CDS isoforms (750 bp to 1750 bp) and increased the abundance of short CDS isoforms (100 bp to 650 bp) (Figures 5D). Thus, the average CDS length of all isoforms decreased after BUD31 silencing (Figure 5E). Moreover, we analyzed the nonsense-mediated mRNA decay (NMD) sensitive splice isoforms of BUD31-regulated AS because coupled AS and NMD might regulate many genes (Jangi and Sharp, 2014). A total of 6,325 NMD-sensitive splice isoforms were identified, and the expression level of the genes that acquired increased NMD-sensitive isoform fraction decreased upon BUD31 knockdown (Figure 5F).

**Figure 5.**
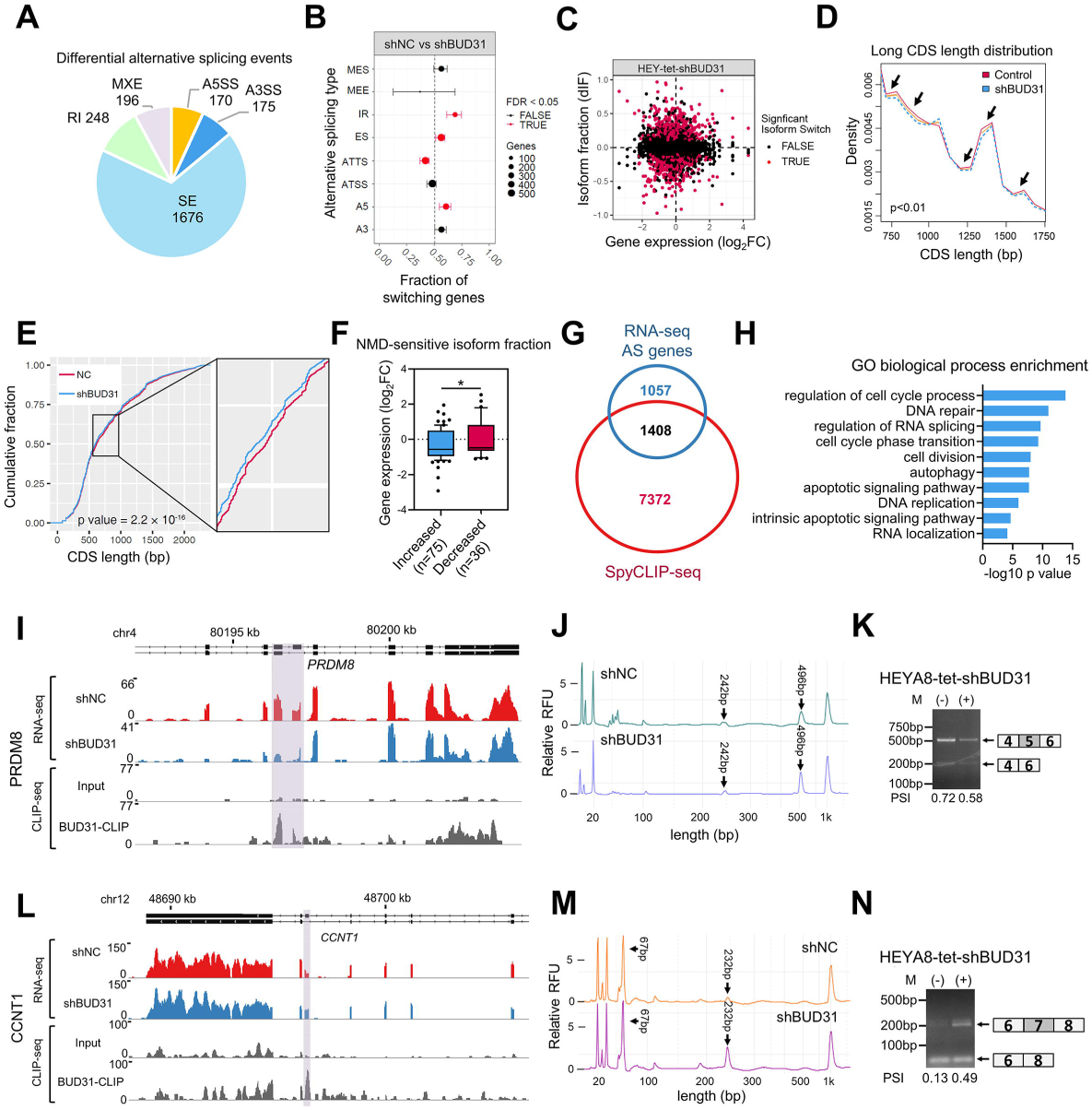
Global identification of AS events regulated by BUD31. (A) Pie chart depicting the proportions of different types of AS events in the RNA-seq data from HEYA8 cells after BUD31 knockdown. Specific differential AS events were analyzed with rMATS and were filtered out with p < 0.05 and |IncLevelDifference| > 0.1. SE, skipped exon; RI, retained intron; A5SS, alternative 5’ splice site; A3SS, alternative 3’ splice site; MXE, mutually exclusive exons. (B) The proportions of genes with different types of AS event changes were analyzed with IsoformSwitchAnalyzeR. FDR < 0.05 was considered statistically significant. (C) The relation between isoform fraction change (dIF) and corresponding gene expression fold change was analyzed. Genes with significant isoform switches are colored in red. (D) Density plot of the long CDS (750 bp to 1750 bp) length distribution in HEYA8 cells with BUD31 knockdown (blue) compared with corresponding controls (red). CDS lengths were calculated using SpliceR with the reconstructed transcripts from cufflinks. The regions reaching statistical significance are shown in orange with a black arrow. (E) Cumulative distribution function plot of CDS lengths of all annotated genes after BUD31 knockdown. The p-value was obtained by the Kolmogorov-Smirnov test. (F) The relation between NMD-sensitive isoform fraction and differential gene expression (p < 0.05). 75 genes with increased NMD-sensitive isoforms fraction (dIF > 10%) after BUD31 knockdown were compared with 36 genes with decreased fraction (dIF < -10%). The log2 transformed gene expression fold changes were shown in boxplot (10-90 percentile), and the p-value was obtained by two-tailed unpaired Student’s t-test. (G) Venn diagram of the 8,780 BUD31-binding genes from the CLIP-seq data and the 2,465 genes that acquired more than one type of AS event in the RNA-seq data. (H) GO biological process enrichment of 1,408 BUD31 candidate targets from (G). (I and L) The AS pattern and BUD31 direct binding sites in *PRDM8* and *CCNT1* were visualized with IGV using the RNA-seq and SpyCLIP-seq data. The gray region highlights the AS region and the BUD31 binding sites. (J and M) Fragment analysis was performed to validate the AS events in *PRDM8* (J) and *CCNT1* (M) in HEYA8 cells with BUD31 knockdown compared with controls. (K and N) Semi-quantitative RT-PCR validated AS events in *PRDM8* (K) and *CCNT1* (N) based on RNA-seq analysis.

In order to identify functional target candidates involved in ovarian cancer progression, we integrated BUD31-bound genes and AS-related genes using CLIP-seq and RNA-seq data. Combined analysis revealed that approximately 57% (1,408/2,465) of the genes with AS events upon BUD31 knockdown were also bound by BUD31 (Figure 5G). Moreover, we performed GO analysis of the 1,408 BUD31 target genes and found they were significantly enriched in apoptosis, cell cycle, splicing, and autophagy (Figure 5H). We subsequently validated several BUD31 target candidates to verify the reliability of the CLIP-seq and RNA-seq data. Seven genes were successfully validated by semi-quantitative RT-PCR and fragment analysis (Figures 5I-5N, 6A-6D). For instance, exon 5 of *PRDM8* was skipped, whereas exon 7 of *CCNT1* was included due to BUD31 ablation (Figures 5I-5N). Collectively, these data imply that BUD31 is a functional regulator of AS and predominantly regulates exon skipping.

### BUD31 promotes *BCL2L12* exon 3 inclusion through direct binding to the pre-mRNA

Of the BUD31-bound and alternatively spliced transcripts, BCL2L12, an anti-apoptotic BCL2 family member, caught our attention. BCL2L12 has been identified as a rational therapeutic target in glioblastoma, which is overexpressed in glioblastoma samples and is low in normal tissues (Jensen et al., 2013). Very recently, therapeutic RNAi targeting *BCL2L12* has been conducted a first-in-human trial in glioblastoma (Kumthekar et al., 2021). We first analyzed RNA-seq data and found BUD31 knockdown promoted exon 3 skipping and resulted in a short isoform (*BCL2L12*-S). Strikingly, BUD31 was shown to bind to exon 3 of *BCL2L12* according to the CLIP-seq and RIP-seq data, indicating that *BCL2L12* is a direct target of BUD31 (Figures 6A and 6B). We next verified the role of BUD31 in the AS of *BCL2L12* by semi-quantitative RT-PCR and fragment analysis. We found that BUD31 knockdown promoted exon 3 skipping to generate increased *BCL2L12*-S expression and decreased level of the full-length isoform (*BCL2L12*-L) (Figures 6C and 6D). We further determined the *BCL2L12*-L/*BCL2L12*-S ratio using isoform-specific primers. Importantly, knockdown of BUD31 in HEYA8 cells significantly decreased the *BCL2L12*-L/*BCL2L12*-S ratio, whereas ectopic expression of BUD31 in A2780 and OVBWZX cells had the opposite effect (Figure 6E). Furthermore, sequence analysis showed that the exon 3 skipping of *BCL2L12* caused a frameshift that introduced a premature termination codon (Figure 6F), and transcripts with such termination codons are predicted to be degraded by NMD (Lindeboom et al., 2016). To verify that *BCL2L12*-S is degraded by NMD, we measured the RNA half-life of *BCL2L12*-S in UPF1 knockdown and control HEYA8 cells treated with the transcription inhibitor actinomycin D. Notably, the half-life of *BCL2L12*-S was significantly increased and the *BCL2L12*-S/ *BCL2L12*-L ratio was higher in UPF1 knockdown cells relative to controls, confirming that *BCL2L12* is sensitive to NMD (Figures 6G). To obtain further evidence for the direct binding of BUD31 to *BCL2L12* pre-mRNA, we performed an RNA immunoprecipitation (RIP) assay in HEYA8 cells with BUD31 antibody or control IgG. RIP-qPCR showed that BUD31 bound to exon 2, exon 3, and intron 3, but not to exon 1 or intron 2 (Figure 6H). An RNA pull-down assay further revealed that BUD31 was more abundantly enriched by wildtype probe, but not by mutant probe spanning exon 3 to intron 3 (Figures 6I). Moreover, an RNA electrophoretic mobility shift assay (EMSA) confirmed the interaction between BUD31 and *BCL2L12* pre-mRNA spanning exon 3 to intron 3 (Figure 6J). These findings strongly support the notion that BUD31 promotes *BCL2L12* AS through direct binding to the pre-mRNA.

**Figure 6.**
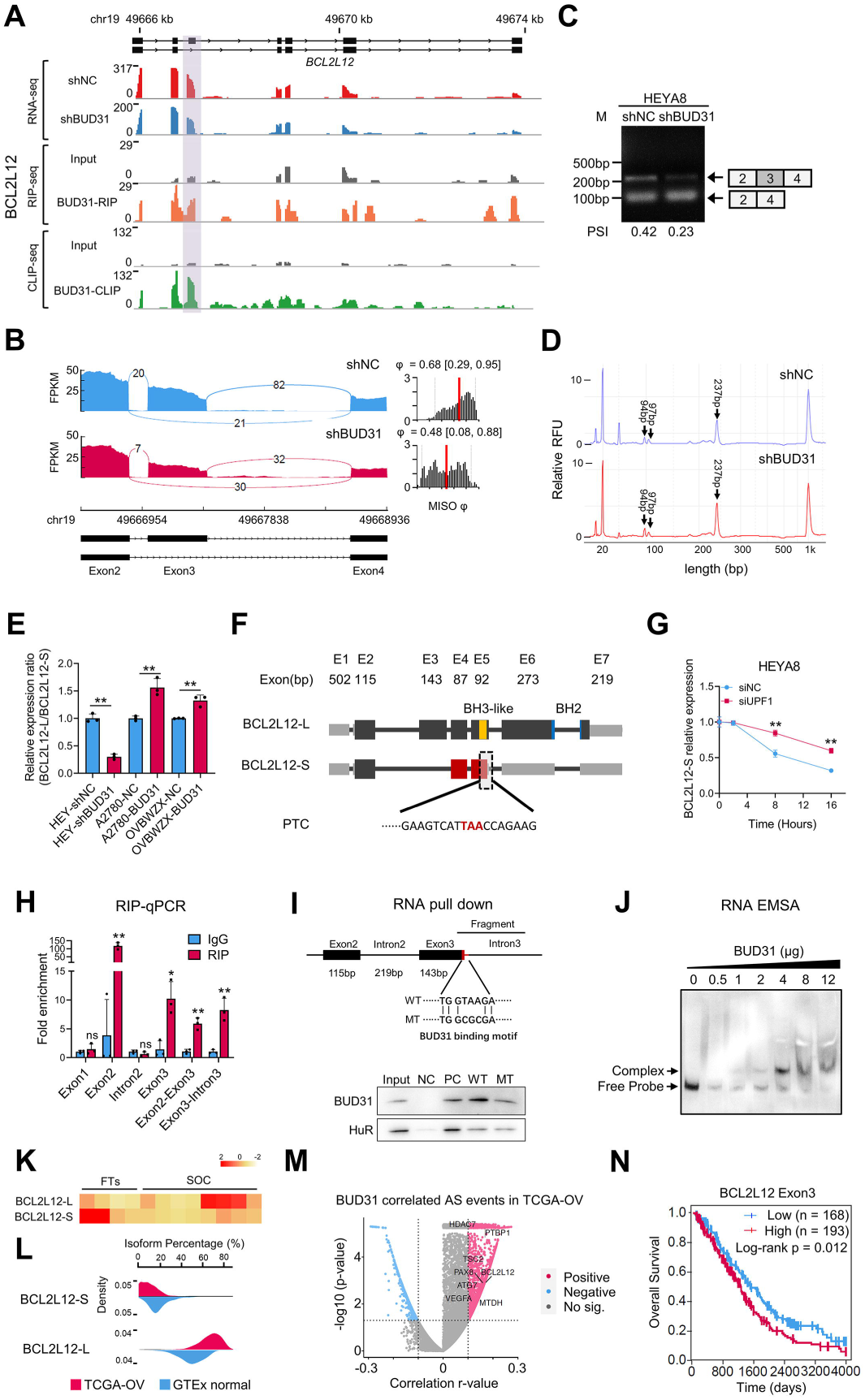
BUD31 promotes *BCL2L12* exon 3 inclusion through direct binding to the pre- mRNA. (A) The AS pattern and BUD31 direct binding sites in *BCL2L12* were visualized with IGV using RNA-seq, RIP-seq, and SpyCLIP-seq data. The gray region highlights the AS region and the BUD31 binding sites. (B) The Sashimi plots of exon 3 skipping in *BCL2L12* in HEYA8 cells with BUD31 knockdown (red) and corresponding controls (blue). The PSI value (φ) for exon 3 skipping was calculated with MISO. (C and D) Semi-quantitative RT-PCR and fragment analysis were performed to validate AS events in *BCL2L12*. (E) The relative expression ratio of *BCL2L12*-L/*BCL2L12*-S was analyzed in ovarian cancer cells with BUD31 knockdown (HEYA8 cells) or BUD31 overexpression (A2780 and OVBWZX cells). (F) Schematic structure of two *BCL2L12* transcripts. *BCL2L12*-L is the full-length transcript, and *BCL2L12*-S is a short transcript lacking exon 3 skipping, which generates a premature termination codon (PTC). (G) *BCL2L12*-S expression was measured by qPCR in UPF1 knockdown and control HEYA8 cells treated with 10 μg/ml actinomycin D at the indicated times. (H) RIP-qPCR was performed to validate the interaction between BUD31 and *BCL2L12* RNA in HEYA8 cells with the anti-BUD31 antibody. (I) RNA pull-down assay showed the interaction between *BCL2L12* RNA and BUD31 protein. (J) RNA EMSA showed the binding of recombinant BUD31 and *BCL2L12* pre-mRNA fragments. The upper band shows the complex of BUD31 protein and *BCL2L12* pre-mRNA. The lower band indicates the free probe. (K) Relative *BCL2L12*-L/S transcript expression was measured by qPCR in SOC samples (n = 8) and FTs (n = 4). (L) Isoform percentage of *BCL2L12*-L and *BCL2L12*-S in SOCs and normal ovaries from the TCGA-OV and GTEx datasets. (M) Correlation between *BUD31* mRNA expression and the PSI value of AS events in the TCGA-OV dataset. P-values and r-values were calculated by Pearson’s correlation (p < 0.05, |r| > 0.1). (N) Kaplan–Meier analyzes the correlation between *BCL2L12* exon 3 expression and overall survival based on TCGA data. The p-values were obtained by two-tailed unpaired Student’s t- test (E, G, H) or log-rank test (N). *p < 0.05, **p < 0.01

To determine the clinical relevance of *BCL2L12* exon 3 skipping in ovarian cancer, we analyzed the expression of *BCL2L12-*L (exon 3 inclusion) and *BCL2L12*-S (exon 3 skipping) in the TCGA ovarian cancer database and found that *BCL2L12*-L was significantly increased whereas *BCL2L12*-S was decreased in SOC (Figures 6K and 6L). In addition, BUD31 was positively correlated with plenty of inclusion events in ovarian cancer, and *BCL2L12* AS was one of them based on PSI value profiles in the TCGASpliceseq database (Figure 6M). More importantly, ovarian cancer patients in the TCGA-OV cohort with exon 3 inclusion showed poor overall survival (Figure 6N), implying that inclusion of *BCL2L12* exon 3 may be involved in ovarian cancer progression.

### BUD31 exerts its oncogenic roles in ovarian cancer by sustaining *BCL2L12* expression

To investigate whether the regulation of *BCL2L12* splicing by BUD31 could be important in mediating the oncogenic role of BUD31 in ovarian cancer, we first analyzed TCGA data and found that BCL2L12 was positively correlated with BUD31 (Figure 7A). Next, we measured the protein level of BCL2L12 in ovarian cancer cells with BUD31 knockdown or overexpression. BUD31 knockdown decreased the protein level of BCL2L12 while forced expression of BUD31 has the opposite effect (Figure 7B). In addition, silencing BUD31 also decreased BCL2L12 expression in the xenograft tumor *in vivo* (Figure S6E). We then conducted rescue experiments to determine whether the phenotype induced by knockdown of BUD31 could be rescued by BCL2L12 overexpression. Indeed, BUD31 knockdown-induced apoptosis was effectively attenuated by BCL2L12 overexpression (Figure 7C and 7D). Additionally, the upregulation of BCL2L12 significantly reversed the inhibition of proliferation and clonogenic ability by BUD31 knockdown in HEYA8 cells (Figures 7E and 7F). Moreover, overexpression of BCL2L12 rescued the inhibition of xenograft growth induced by silencing BUD31 (Figures 7G, 7H and 7I). Especially, BCL2L12 was overexpressed in SOCs, and high BCL2L12 expression levels predicted a poor prognosis for patients with SOC (Figures 7J, 7K and 7L). These results suggest that BUD31 exerts its oncogenic roles in ovarian cancer partially through regulating *BCL2L12*.

**Figure 7.**
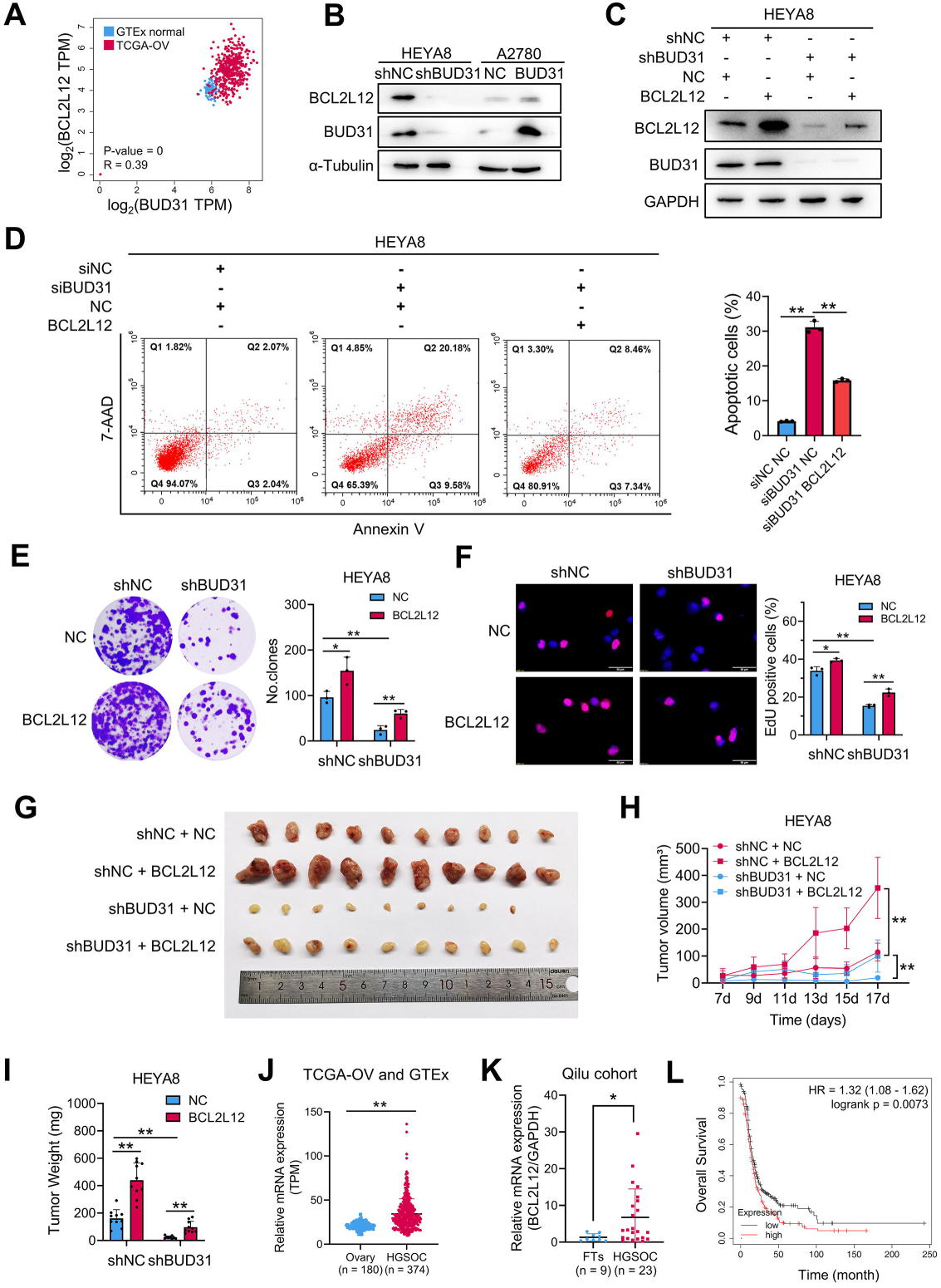
BUD31 exerts its oncogenic effects in ovarian cancer by sustaining *BCL2L12* expression. (A) Correlation analysis of mRNA expression between *BUD31* and *BCL2L12* in SOCs and normal ovaries from the TCGA-OV and GTEx datasets. (Pearson’s r = 0.39). (B) Western blot analysis of BCL2L12 protein expression in HEYA8 cells with BUD31 knockdown and in A2780 cells with BUD31 overexpression. (C) Immunoblot analysis of BUD31 and BCL2L12 protein levels in HEYA8 cells transfected with shBUD31 or BCL2L12 overexpression vector. (D-F) Apoptosis (D), clonogenic (E), and EdU (F) assays for investigating the potential of BCL2L12 to rescue the loss of BUD31 in terms of apoptosis and proliferation. (G-I) Xenograft experiments by subcutaneous injection were conducted in HEYA8 cells with dox-inducible BUD31 knockdown or BCL2L12 overexpression vector. Representative image (G), volume curves (H), and weight (I) of xenograft tumors showed that BCL2L12 could partially rescue the inhibitory effect of BUD31 on tumor growth (n = 10). (J) BCL2L12 expression was analyzed in SOCs from TCGA-OV (n = 374) and in normal ovaries from GTEx (n = 180). (K) *BCL2L12* mRNA expression was determined by qPCR in SOC (n = 23) and FT (n = 9) samples. (L) Kaplan–Meier analysis of BCL2L12 expression on the overall survival of ovarian cancer patients based on cohorts from Kaplan–Meier Plotter. The high and low expression groups were separated based on the autoselect cutoff. All functional experiments were performed with n = 3 biological repeats. The p-values were determined by a two-tailed unpaired Student’s t-test (D, E, F, H, I, J, K), or log-rank test (L). *p < 0.05, **p < 0.01

### ASO-mediated *BCL2L12* exon skipping induces apoptosis of ovarian cancer cells

Splice-switch ASOs are a promising strategy for the treatment of various diseases, including cancer, and ASOs have been approved for the treatment of spinal muscular atrophy and Duchenne muscular dystrophy (Scharner et al., 2019). To identify effective ASOs that promote *BCL2L12* exon 3 skipping, we analyzed the BUD31 binding region on *BCL2L12* based on CLIP-seq and designed three ASOs modified with phosphorothioate linkages (Figure 8A). We examined the endogenous *BCL2L12* splicing in A2780 cells transfected with ASOs and controls and found that ASO2 and ASO3 efficiently reduced *BCL2L12*-L (exon 3 inclusion) and increased *BCL2L12*-S (exon 3 skipping) levels (Figure 8B). Consistent with this, BCL2L12 protein expression was significantly decreased upon ASO2 and ASO3 treatment (Figures 8C). ASO2 was chosen for subsequent experiments because it had the strongest effect on exon 3 skipping of *BCL2L12*. To further determine the effect of ASO2 on the splicing switch of *BCL2L12*, ASO2 was transfected into HEYA8 and A2780 cells and splicing of exon 3 was measured by RT-PCR. A dose-dependent increase in exon 3 skipping was observed after ASO2 treatment (Figure 8D), and ASO2 was more potent in A2780 cells (half-maximal effective concentration [EC50] of 52.91 nM) than in HEYA8 cells (EC50 of 63.77 nM) (Figure 8E). ASO2 decreased BCL2L12 expression in a dose and time-dependent manner (Figures 8F and 8G). Importantly, ASO2 treatment induced apoptosis of A2780 and HEYA8 cells as determined by flow cytometry and cleaved-caspase3 expression (Figures 8F- 8H). Meanwhile, the EdU incorporation assay showed that ASO2 significantly suppressed the proliferation of ovarian cancer cells (Figure 8I). We further determined the half-maximal inhibitory concentration (IC50) values for ASO2 to be 74.27 nM in A2780 cells and 73.70 nM in HEYA8 cells (Figure 8J). Moreover, a subcutaneous xenograft model was established using HEYA8 cells and showed that ASO2 treatment significantly reduced the tumor size and increased apoptosis (Figure 8K-8N). Taken together, these results suggest that ASO2 inhibits ovarian cancer cells proliferation by regulating *BCL2L12* exon 3 skipping.

**Figure 8.**
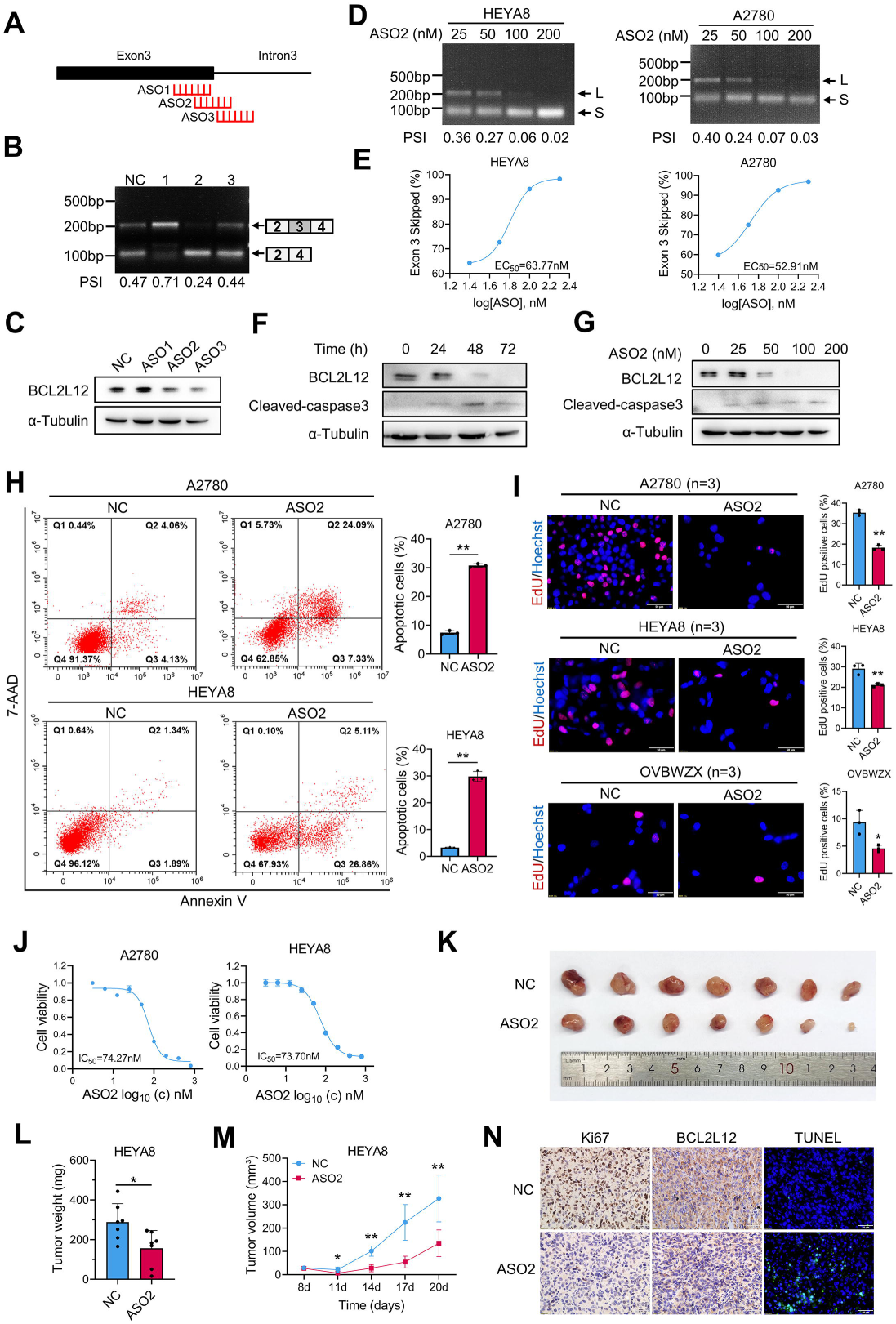
ASO-mediated *BCL2L12* exon skipping induces apoptosis of ovarian cancer cells. (A) Schematic diagram of ASO target sites on *BCL2L12* based on the BUD31 binding region on *BCL2L12* as determined from the CLIP-seq data. (B) RT-PCR analysis of the *BCL2L12* AS pattern in response to ASOs. (C) Western blot analysis of the BCL2L12 protein level in A2780 cells transfected with ASOs. (D) Semi-quantitative RT-PCR analysis of the *BCL2L12* AS pattern in HEYA8 and A2780 cells transfected with ASO2. (E) Quantification of the bands shown in (D). Dose-dependence curve of ASO2-treated HEYA8 and A2780 cells showing increased skipping of exon 3 [(exon 3 skipped/exon 3 skipped + full-length) × 100] in relation to the log of the dose. The EC_50_ was calculated in HEYA8 (63.77 nM) and A2780 (52.91 nM) cells. (F-G) BCL2L12 and cleaved-caspase3 were measured by Western blot in A2780 cells treated (F) with 200 nM ASO2 for different times (0, 24, 48, and 72 h) and (G) with different concentrations (0, 25, 50, 100, 200 nM) of ASO2 for 72 h. (H) Apoptotic cells were detected by flow cytometry after staining with Annexin V/7-AAD in A2780 and HEYA8 cells treated with ASO2 (200 nM). (I and J) EdU (I) and MTT (J) assays were performed in ovarian cancer cell lines treated with 200 nM ASO2. The IC_50_ was calculated based on the MTT assay. (K-M) ASO2 intratumoral injection to subcutaneous tumor xenografts using HEYA8 cells. The xenograft model showed the inhibitory effect of ASO2 on tumor growth (n = 6 mice per group) (K). The tumor weight (L) and volume (M) were measured for each group. The BCL2L12 and Ki67 expression levels were evaluated with immunohistochemical staining, and the apoptosis level was determined by a TUNEL assay (N). All functional experiments were conducted with n = 3 biological repeats. The p-values were obtained by two-tailed unpaired Student’s t-test (H, I, L, M). *p < 0.05, **p < 0.01.

## DISCUSSION

Dysregulated expression of splicing factors and perturbed splicing have been shown to drive carcinogenesis and tumor development. We carried out a screen for survival-related splicing factors in SOC using TCGA data. We found that BUD31 was commonly overexpressed in SOC and that a high level of BUD31 was associated with poor prognosis, and pan-cancer analysis showed that BUD31 was overexpressed in various cancer types. We next investigated the functional role of BUD31 and found that BUD31 promoted the proliferation and survival of ovarian cancer cells and xenograft tumor growth. In breast cancer cells, BUD31 is required for cell migration (Koedoot et al., 2019). These findings suggest that BUD31 has oncogenic potential and is closely related to the unfavorable prognosis of patients with ovarian cancer.

Synthetic lethality provides a promising strategy for cancer therapy (O’Neil et al., 2017). PARP inhibitors are the first clinically approved drugs based on their synthetic lethality in BRCA1/2 mutant ovarian cancer (Mateo et al., 2019). In MYC activated state, *BUD31* has been identified as a MYC-synthetic lethal gene, BUD31 in required for spliceosome assembly and catalytic activity. Depletion of BUD31 in MYC-hyperactive cells leads to global intron retention (Hsu *et al*., 2015). However, the BUD31-regulated cancer-specific splicing events and its binding motif remain largely unknown. For the first time, we performed CLIP-seq to map genome-wide BUD31-RNA interactions. We found BUD31 binding sites were highly enriched in exon-intron regions near both the 3’ and 5’ splicing sites. We further revealed that BUD31 inhibition results in extensive exon skipping and decreased abundance of long CDS isoforms. We also identified four BUD31-binding RNA motifs using the HOMER algorithm. In the combined analysis of CLIP-seq and RNA-seq data, we identified multiple BUD31 potential direct binding targets. These findings reveal that BUD31 globally regulates AS through direct binding its RNA targets.

Furthermore, we provide strong evidence that *BCL2L12*, a member of the Bcl-2 anti-apoptotic family (Bcl2-like-12), is a crucial AS target of BUD31. BCL2L12 has been identified as a therapeutic target in glioblastoma (Jensen *et al*., 2013). RNA interference-based spherical nucleic acids targeting *BCL2L12* has been conducted phase 0 first-in-human trial in glioblastoma (Kumthekar *et al*., 2021). BCL2L12 expression is upregulated in human glioblastomas and confers resistance to apoptosis (Stegh et al., 2007), and BCL2L12 impedes p53-dependent DNA damage-induced apoptosis through direct interaction with p53 (Stegh et al., 2010). BCL2L12 induces Th2 polarization and contributes to inflammation in the intestinal mucosa (Li et al., 2018). Importantly, we found that BUD31 knockdown promoted exon 3 skipping to generate an increased expression of *BCL2L12*-S and decreased expression of the full-length isoform *BCL2L12*-L. Interestingly, *BCL2L12*-S was then degraded by NMD, and the cells subsequently underwent apoptosis. BUD31 was shown to bind to exon 3 of *BCL2L12* according to the CLIP-seq and RIP-seq data. BUD31 protein and *BCL2L12* pre-mRNA interactions were verified by EMSA and RNA pull-down assays. Intriguingly, BUD31 exerts its oncogenic roles in ovarian cancer by regulating BCL2L12. More importantly, *BCL2L12*-L was significantly increased in SOCs and higher levels were correlated with poor overall survival. Thus, our work supports the hypothesis that BUD31 stimulates the inclusion of exon 3 to generate *BCL2L12*-L and promotes ovarian cancer progression.

RNA-based therapeutics is coming of age. SARS-CoV-2 mRNA-based vaccines are approximately 95% effective in preventing COVID-19 (Turner et al., 2021). Several ASOs have been approved for the treatment of spinal muscular atrophy and Duchenne muscular dystrophy (Scharner *et al*., 2019). Spinraza is an FDA-approved drug that can modulate splicing of the *SMN2* gene to generate full-length SMN2 protein and thus improve the motor function of spinal muscular atrophy patients (Wan and Dreyfuss, 2017). PKM2 is an isoform of the *PKM* gene and plays a crucial role in the Warburg effect in cancer, and splice switching from PKM2 to PKM1 by ASOs restores the sensitivity of cancer cells to chemotherapy (Calabretta et al., 2016). ASO-mediated exon 6 skipping of *MDM4* reduces the amount of full-length MDM4 and inhibits melanoma cell growth (Dewaele et al., 2016). Finally, ASOs have been shown to induce the redirection from Bcl-xL to Bcl-xS and thus to induce apoptosis of melanoma cells in vitro and to inhibit xenograft tumor burden in vivo (Bauman *et al*., 2010). Here, we designed ASOs to target *BCL2L12* exon 3 near the 5′ splice sites, which results in exon 3 skipping. We then examined the endogenous *BCL2L12* splicing in A2780 cells transfected with ASOs and found that ASO2 efficiently reduced *BCL2L12*-L levels and increased *BCL2L12*-S levels. Consistent with this, BCL2L12 protein expression was significantly decreased upon ASO2 treatment. Importantly, ASO2 was able to induce apoptosis and to inhibit xenograft tumors of ovarian cancer cells both in vitro and in vivo. Our findings thus suggest that ASO-mediated *BCL2L12* exon 3 skipping is a promising strategy for cancer therapy.

In conclusion, our study suggests that BUD31 acts as an oncogenic splicing factor and prognostic marker in ovarian cancer. We further identified the binding motif and the preferred genome-wide binding pattern of BUD31 by crosslinking immunoprecipitation sequencing analysis. Specifically, BUD31 overexpression drives an oncogenic splicing switch in *BCL2L12* to produce *BCL2L12*-L that in turn increases the survival and proliferation of ovarian cancer cells. Inhibition of BUD31 or the use of ASOs may provide novel therapeutic strategies for ovarian cancer.

## MATERIAL AND METHODS

### Nude mouse xenograft model

In the subcutaneous xenograft model, female BALB/c-nude mice (6–8 weeks old) were randomly divided into two groups (4–8 mice per group) and injected subcutaneously with doxycycline-induced BUD31 knockdown HEYA8 cells or BUD31 overexpression ID8 cells as previously described (Wang et al., 2017). Doxycycline (1.2 g/L) mixed with 5% sucrose was fed to the experimental group, while 5% sucrose alone was fed to the control group. ASO (Tsingke) intratumor injection was applied after the subcutaneous tumor reached 5mm in diameter. 5 nmol ASO mixed with 3 μl lipo2000 in 25 μl Opti-MEM was administered every 3 days. When the study finished, the mice were anesthetized, and the tumor volume and weight were measured.

In the living image xenograft model, female BALB/c-nude mice (6–8 weeks old) were randomly divided into two groups (6 mice per group) and injected intraperitoneally with luciferase-expressing ovarian cancer HEYA8 cells with a dox-inducible BUD31 knockdown system. Administration of doxycycline (1.2 g/L) in the drinking water started one week after the cell implantation. Three weeks later, the mice were anesthetized with 4% sterile chloral hydrate (7–10 µl/g body weight, Sangon) and D-Luciferin sodium salt (15 mg/ml, dissolved in DPBS, 150 µg/g body weight, Yeason) was injected intraperitoneally. Bioluminescence images were captured 10 min later using an imaging system (PerkinElmer). The Shandong University Animal Ethics Research Board approved the animal experiment procedures (SDULCLL2019-2-08).

### Human tissue samples

SOC specimens and FTs were collected from the Department of Obstetrics and Gynecology, Qilu Hospital, Shandong University, from April 2009 to July 2015 as previously described (Wang *et al*., 2021). The SOC samples were obtained from patients with primary ovarian cancer who had not undergone any previous surgery or chemotherapy. In addition, FTs were obtained from patients who underwent total hysterectomy and bilateral salpingo-oophorectomy for uterine diseases or benign neoplastic adnexal pathological changes. Fresh tissue samples were collected within 2 h of surgery and were sliced to 5 mm^3^ and immersed in 10 vol of RNALater (Ambion, Austin, TX). The tissue samples were stored at –80 °C. All patients provided informed consent, and ethical approval was granted by the Ethics Committee of Shandong University (SDULCLL2019-1-09).

### Cell lines

Human ovarian cancer cell lines A2780 and HEYA8 were obtained from the Jian-Jun Wei lab, Northwestern University. OV90 and OVCAR3 cell lines were purchased from the American Type Culture Collection. The mouse ovarian surface epithelial cell line ID8 was purchased from Sigma-Aldrich (SCC145). Cell lines were validated by STR profiling. A2780, HEYA8, ID8, and OV90 cells were cultured in Dulbecco’s modified Eagle’s medium (DMEM) (Gibco, Invitrogen) containing 10% fetal bovine serum (FBS) (Gibco) and 1% penicillin/streptomycin (Macgene). OVCAR3 cells were cultured in RPMI 1640 (Macgene) supplemented with 20% FBS, 1% penicillin/streptomycin, 1% sodium pyruvate (Sigma), 0.3% glucose (Corning), and 10 ng/mL insulin (Sigma). The cells were cultured at 37°C in a humidified incubator containing 5% CO_2_.

### Primary culture of ascites-derived ovarian cancer cells

Patient ascites were collected from the Department of Obstetrics and Gynecology, Qilu Hospital, Shandong University, upon the patient’s informed consent. Ascites-derived ovarian cancer cells OVBWZX were obtained from a 54 years old female patient diagnosed with high-grade serous ovarian cancer by clinical pathology before receiving chemotherapy and surgical therapy. Primary ovarian cancer cells were cultured according to the previous studies (Shepherd et al., 2006; Thériault et al., 2013). Briefly, after receiving freshly isolated fluid in a sterile vacuum container, 25 ml of ascites fluid was mixed with an equal volume of MCDB/M199 medium in T-75 flasks. Cells were incubated for 3–4 days prior to the first change of complete medium. The medium was then changed every 2–3 days until the flasks were confluent and cells were passaged at a 1:2 dilution. Experiments were performed using cells at passage 2 through passage 6. Primary ovarian cancer cells were separated into 30 divisions and frozen in 70% v/v MCDB/M199 medium, 20% v/v FBS, and 10% dimethylsulfoxide (DMSO).

### Lentiviral infection and RNA interference

The BUD31-pENTER overexpression plasmid and the BCL2L12-pENTER plasmid were from WZ Biosciences. The full-length open reading frame of *BUD31* was cloned with a ClonExpress®II One Step Cloning Kit (Vazyme) into the modified doxycycline-inducible lentiviral vector pTRIPZ (Zhao *et al*., 2019), which was a gift from Ligang Wu’s lab, University of Chinese Academy of Sciences. The shBUD31 sequence targeting BUD31 was cloned into the pZIP-TRE3G plasmid (Transomic). The lentivirus vectors were transfected into HEK293T cells together with psPAX2 and pMD2.G to produce lentivirus particles. Stable cell lines were established by lentivirus infection followed by puromycin (2 μg/ml) selection for 2 weeks. RiboBio prepared a GenOFF st-h-BUD31 suite containing three siRNAs and GenOFF st-h-BCL2L12. Plasmids and siRNAs were transfected into cells using Lipofectamine 2000 reagent (Thermo Fisher Scientific) following the manufacturer’s instructions. RNA and protein were extracted 48 h or 72 h after transfection or doxycycline induction.

### Recombinant protein expression and purification

*Escherichia coli* BL21(DE3) competent cells (Vazyme) were transformed with SpyCatcher expression plasmid, a gift from Ligang Wu’s lab, and the BUD31 expression pET-28a plasmid. A single colony was picked and inoculated into 1 ml Luria-Bertani medium containing 0.5 mg/ml kanamycin (BBI Life Science, Shanghai, China) for 10 h at 37°C. A total of 100 µl cells were inoculated and cultured at 37°C until the absorbance at 600 nm reached 0.4–0.6, and this was followed by induction with 1 mM isopropyl β-D-thiogalactoside (IPTG) (BBI, Shanghai, China) at 30°C for 6 h. Cells were harvested and lysed using an ultrasonic cell crusher (SCIENTZ, Ningbo, China). The lysates were centrifuged at 12,000 rpm for 50 min, and the soluble fraction was purified with a His-tag Protein Purification Kit (Beyotime, China) according to the manufacturer’s protocol. The eluates were dialyzed overnight against dialysis buffer (20 mM Tris-HCl pH 7.8, 150 mM NaCl, 10% glycerol, and 1 mM DTT). SDS-PAGE and Coomassie blue staining checked the purity of each fraction. Purified proteins were aliquoted and stored at –80°C.

### RNA isolation and PCR analysis

According to the manufacturer’s instructions, total RNA was extracted from cultured cells or fresh tissues with a Cell Total RNA Isolation Kit (Foregene). Total RNA was reverse transcribed into cDNAs using HiScript II Q RT SuperMix for qPCR (Vazyme), and qPCR was performed with SYBR Green mix (Vazyme) and the Applied Biosystems QuantStudio 3. GAPDH served as the endogenous control. The 2^−ΔΔCT^ method was used for the relative quantification of the qPCR data. Semi-quantitative RT-PCR and Qsep100 Bio-Fragment Analyzer (BiOptic) were used to analyze alternative spliced products. Primer sequences were designed for the constitutively expressed flanking exons (Ferre, 1992), and 2 × Taq Master Mix (Dye Plus) (Vazyme, P112-01) was used to simultaneously amplify isoforms that included or skipped the target exon. Primer sequences are listed in Table S5.

### Western blot

The samples were lysed on ice in Western and IP Cell Lysis Buffer, and the protein concentration was determined using the bicinchoninic acid protein assay (Beyotime). SDS-PAGE was used to separate protein samples. The membrane was blocked with 5% skim milk before overnight incubation with primary antibodies at 4°C. Horseradish peroxidase (HRP)- conjugated secondary antibodies and an electrochemiluminescence system (GE Healthcare, UK) were used to detect specific proteins. All antibody information is listed in the Table S6.

### IP-MS

HEYA8 cells with endogenously expressed BUD31 were harvested, and cell pellets were lysed on ice with Western and IP Cell Lysis Buffer (Beyotime Institute of Biotechnology). Whole-cell extracts were incubated with 5 μg BUD31 antibody (Proteintech) overnight for 1 h at 4°C, then incubated with magnetic Protein A/G beads (Millipore) for 2 h at 4°C. Beads were washed with Western and IP Cell Lysis Buffer three times, and the immunocomplex was resuspended in 1× SDS-PAGE loading buffer and separated followed by Coomassie brilliant blue staining. LC-MS/MS was conducted by PTM-Biolab using a Thermo Fisher LTQ Obitrap ETD. The peptides were confirmed by western blot. The data of the IP-MS are given in Table S3.

### Cell proliferation assays

Cell viability and proliferation were determined with a methyl-thiazolyl diphenyl-tetrazolium bromide (MTT) assay. Cells were seeded in 96-well plates at densities of 1 × 10^3^ cells per well. After culturing for the designated time, 10 μl of MTT (5 mg/ml) was added to each well and incubated for 4 h at 37 °C. Cell growth was monitored over the following 5 days, and the IC50 was determined 48 h after treatment. The supernatant was discarded after centrifugation, and 100 μl of dimethylsulfoxide was added to each well and the absorbance was measured at 570 nm using a microplate reader (Bio-Rad, Hercules, CA, USA). According to the manufacturer’s instructions, an EdU cell proliferation assay was performed using a Cell-Light EdU Apollo567 In Vitro Kit (RiboBio, Guangzhou, China). Briefly, cells were seeded on glass coverslips in 24-well plates at densities of 3–4 × 10^4^ cells per well and then incubated with the cell culture medium containing EdU for 20–30 min. The cells were then fixed and stained with Apollo567 fluorescent dye and Hoechst 33342.

### Clonogenic assay

Cells were seeded in a six-well plate (1,000–2,000 cells per well) and cultured for 1–2 weeks. Colonies were fixed with methanol and stained with 0.1% crystal violet (Solarbio), and the number of colonies was counted with Image J. The data are presented as the mean ± SD of three independent experiments.

### Flow cytometry and TUNEL assays

Flow cytometry analysis for cell apoptosis was performed using an Annexin V-PE/7-AAD Apoptosis Detection Kit (Vazyme, A213-01). In H_2_O_2_-induced apoptosis experiments, cells were treated with H_2_O_2_ with a final concentration of 400 μM for 4 h before apoptosis detection. Cells were digested with ethylenediaminetetraacetic acid (EDTA)-free trypsin (Macgene, CC035) for 3 min, collected by centrifugation, washed with ice-cold phosphate-buffered saline (PBS) and resuspended at a density of 5 × 10^5^ cells/ml with 100 µl 1× Binding Buffer. Then 5 μl Annexin V-PE and 5 μl 7-AAD were added and incubated for 10 min in the dark. Finally, cells were incubated with an additional 400 µl 1× Binding Buffer and analyzed within 20 min by CytoFLEX S (Beckman Coulter Life Science). At least 1 × 10^4^ cells were analyzed to determine the percentage of apoptotic cells. For TUNEL assays, the TUNEL Cell Apoptosis Detection Kit (KEYGEN, KGA703) was used according to the manufacturer’s protocol.

### IHC staining of tissue sections

Formalin-fixed and paraffin-embedded tissues or tissue microarray sections were deparaffinized in xylene and rehydrated with a graded series of ethanol solutions. Antigen retrieval was performed in EDTA by heating in a microwave. Tissue slides were blocked with 1.5% normal goat serum and incubated with primary antibodies against Ki-67 (1:200 dilution, CST, 9129S) and BUD31 (1:250 dilution, Proteintech, 11798-1-AP) overnight at 4°C. The sections were then incubated with the secondary antibody and stained with diaminobenzidine (DAB). The final IHC staining score for the tissue microarray was determined as we described previously (Wang *et al*., 2021). Specifically, high (Score ≥ 7) and low (Score < 7) expression of each sample was determined by two pathologists based on the intensity and extent of staining across the tissue microarray section. The clinical information and expression score are listed in Table S7.

### Immunofluorescence staining

HEYA8 and OVBWZX cells were fixed with 4% paraformaldehyde for 30 min at room temperature, followed by permeabilization with 0.5% Triton X-100 in PBS for 15 min at room temperature. Tissue slides were processed as described above for immunohistochemistry staining. Samples were then blocked with 1% BSA for 1 h at room temperature and incubated with primary antibodies against SC35 (1:500 dilution, Abcam, ab204916), BUD31 (1:250 dilution, Proteintech, 11798-1-AP), and α-Tubulin (1:400 dilution, Proteintech, 66031-1-Ig) overnight at 4°C and then incubated with secondary antibody for 1 h at room temperature in the dark. The images were captured on an Andor Revolution confocal microscope system or an Olympus BX53 microscope system.

### RNA immunoprecipitation (RIP) assay

HEYA8 cells were collected for the RIP assay, which was performed using the EZ-Nuclear RIP Kit (17-701, Merck Millipore) following the manufacturer’s instructions. In brief, cells were collected and lysed in RIP lysis buffer, and the lysates were incubated with magnetic beads coated with anti-BUD31 antibody (Proteintech, 11798-1-AP) at 4°C overnight. The beads combined with immunocomplexes were washed with RIP wash buffer six times and digested by protease K, and RNA was extracted with phenol/chloroform/isoamyl alcohol (125:24:1 mixture). Both input and RIP samples were prepared for next-generation sequencing by Ribobio Biotechnology Company.

### SpyCLIP assay

SpyCLIP was performed according to a previous study (Zhao *et al*., 2019) with several modifications. Briefly, HEYA8 cell lines transfected with doxycycline-inducible SpyTag- FLAG-BUD31 expression lentivirus were induced with doxycycline for 72 h before harvesting. Cells were crosslinked and irradiated at 400 mJ/cm^2^ in a UV Crosslinker (UVP, CL-1000). Cell nuclei were isolated with a Nucleoprotein Extraction Kit (Sangon) before lysis. Turbo DNase (2 U/μl, Invitrogen, AM2238) and 1:200 diluted RNase I (100 U/μl, Invitrogen, AM2295) were used to remove the DNA and to fragment the RNA. The mixed lysate was incubated with anti-FLAG magnetic beads (MBL, M185-11) at 25°C for 40 min. After removing RNA-binding proteins from the FLAG beads with phosphoserine phosphatase, the mixture was incubated with fresh pre-washed SpyCatcher beads at 25°C for 1 h. Stringent washes were conducted according to the manufacturer’s instructions, and the beads were digested with proteinase K (Roche, 3115828001). RNA was purified and concentrated with the Spin Column RNA Cleanup & Concentration Kit (Sangon), and the sequencing library was constructed with the NEBNext Ultra RNA Library Prep Kit for Illumina. High-throughput sequencing of the SpyCLIP libraries was performed on a HiSeq 2500 using the PE150 sequencing strategy (Novogene).

### RNA pull-down assay

*BCL2L12* pre-mRNA fragments were cloned from human placenta genomic DNA. Primers used for generating wildtype and mutant *BCL2L12* fragments and the target sequence are listed in Table S5. With the constructed T7 promoter ahead of the target sequence, the RNAMAX-T7 in vitro transcription kit (RiboBio) was used to transcribe the RNA of the *BCL2L12* fragment. The fragment was labeled with biotin using the Pierce™ RNA 3′ End Desthiobiotinylation Kit (Thermo Fisher Scientific), and RNA pull-down was performed using the Magnetic RNA Protein Pull-Down Kit (Thermo Fisher Scientific) according to the manufacturer’s protocol. The negative control was poly(A)_25_ RNA, and the positive control was the 3’ untranslated region of the androgen receptor RNA. The proteins were detected by western blot analysis.

### RNA EMSA

The RNA EMSA was performed with a CoolShift-BTr RNA EMSA Kit (Viagene) according to the manufacturer’s instructions with modifications. Briefly, 10 ng biotin-labeled RNA probe per sample was first heated at 85°C for 3 min to relax secondary structures. The probe was then incubated with recombinant His-tagged BUD31 at different concentrations for 40 min, and 5% nondenaturing polyacrylamide gel electrophoresis was conducted in 0.25× cool Tris-borate-EDTA (TBE) buffer at 120V for 70 min. RNA-protein complexes were transferred to the membrane in 0.5× TBE at 390 mA for 40 min. After immobilization and crosslinking with 600 mJ UV in a CL-1000 UV linker, the membrane was blocked and conjugated with HRP. The chemiluminescence was detected with an ECL system (GE Healthcare, Little Chalfont, Buckinghamshire, UK).

### RNA-seq and quantification

Total RNA was isolated from BUD31 knockdown and control HEYA8 cells (three biological replications of each sample) using TRIzol reagent (Invitrogen) according to the manufacturer’s protocol. Poly(A) sequencing libraries were prepared using the Illumina TruSeq-stranded-mRNA protocol after RNA quality was assessed using an Agilent Technologies 2100 Bioanalyzer with the application of an RNA integrity number > 7.0. Adenylated mRNAs were isolated using oligo-d(T) magnetic beads (two rounds). The RNA-seq library was paired-end sequenced using Illumina HiSeq 4000 at LC-Bio. After obtaining paired-end reads, clean reads were aligned to the hg38 genome with HISAT2 (version 2.2.0) and sorted with samtools (version 1.9). Mapped reads were visualized with the Integrative Genomics Viewer (IGV). Transcripts were reconstructed with StringTie (1.3.0), and differential expression was analyzed with edgeR. The cutoff was set as q < 0.05 and fold change (FC) > 1.7 or < 0.6.

### AS and isoform switch analysis

The mapped reads aligned by HISAT2 were used for further analysis, and AS events were identified mainly by rMATS (version 4.1.0) (Shen et al., 2014). HEYA8 cells with BUD31 knockdown and corresponding controls (n = 3 biological repeats) were used in the analysis. AS events were classified into skipped exon, retained intron, alternative 5’ splice site, alternative 3’ splice site, and mutually exclusive exons. Significant events were filtered out with p < 0.05 and |IncLevelDifference| > 0.1. AS events in each sample were also identified with ASprofile (Florea et al., 2013). MISO (Mixture of Isoforms, version 0.5.4) (Katz et al., 2010) analysis was further conducted to confirm the results. Considering the index version of MISO, clean reads were remapped to human genome hg19 with HISAT2. The MISO parameters were --read-len 141 --paired-end 240 117, and *BCL2L12* exon 3 skipping was plotted with the sashimi plot program. Global AS analysis was performed as part of the IsoformSwitchAnalyzeR results.

For isoform switch analysis, transcript expression was determined with Salmon (version 0.6.0) (Patro et al., 2017) using the quant function with -l U -p 8. The quantification results were imported into R for further analysis. Global isoform fractions and AS events were analyzed with IsoformSwitchAnalyzeR (version 3.13) (Vitting-Seerup and Sandelin, 2019), an R package stored in Bioconductor. In addition, the coding potentials of the transcripts were determined with the Coding Potential Assessment Tool (version 1.2.1) (Wang et al., 2013). Domain information was annotated with Pfam (version 34.0) (Mistry et al., 2021), and Signalp (version 5.0b) (Almagro Armenteros et al., 2019) was used for signal peptide analysis.

### CDS and UTR length analysis

The elements length of each transcript was determined with SpliceR (version 1.14.0) (Vitting-Seerup et al., 2014) using the transcripts reconstructed with Cufflink. The enrichment of each transcript was calculated by Cufflink. Length distribution was visualized with a density plot, and statistical differences were calculated inside the R package sm (version 2.2). Cumulative distribution was analyzed by the Kolmogorov-Smirnov test and plotted by ggplot2. NMD sensitivity was determined with SpliceR. For a transcript to be marked as NMD-sensitive, the minimum distance from a stop codon to the final exon-exon junction was 50 nucleotides.

### Identification of SpyCLIP crosslinking sites and binding motif analysis

Crosslinking sites were identified using the previously described iCLIP analysis pipeline (Busch et al., 2020). The adaptor sequence and low-quality reads were removed with TrimGalore (version 0.6.1), and the quality of the clean reads was checked with FastQC (version 0.11.9). rRNA sequences were removed with bowtie (version 1.2.3) (Langmead et al., 2009). The remaining reads were mapped to the human genome (hg38) using the STAR software (version 2.7.1a) (Dobin et al., 2013). PCR duplicates of uniquely mapped reads were removed using Picard (version 2.25.5) with MarkDuplicates. The remaining reads were considered usable reads for identifying crosslinking sites. Mapped reads were visualized in IGV (Robinson et al., 2017). Two technical replicates of the SpyCLIP samples were merged for PURECLIP (version 1.2.0) (Krakau *et al*., 2017) analysis with -ld -nt 16 -dm 80. Individual crosslink sites (< 80 nt) were merged as raw binding regions. Binding regions that acquired more than three crosslink sites were used for further analysis as previously suggested (Busch *et al*., 2020), and these regions were annotated using HOMER (version 4.11) (Duttke et al., 2019) into exon, intron, promoter, intergenic, UTR’3, UTR’5, etc. The 80-nt regions around the center of each binding region were extracted and used to identify the de novo BUD31 binding motif using HOMER’s findMotifs program (-len 6,8,10,12 -S 10 -rna -p 4). Motifs were matched to the genome position with scanMotifGenomeWide.pl and visualized with Deeptools.

### Visualization of the binding region distributions around the regulated exons

The RNA-seq data upon knockdown of BUD31 in HEYA8 cells were obtained as described above. Alternative splicing events were identified by rMATS, and 2,000 randomly chosen exons were used as controls. The enrichment of the BUD31 binding signal near the regulated exons was analyzed by deeptools (version 3.1.3) and was calculated using the same code as previously reported (Chakrabarti et al., 2017).

### Data source, functional enrichment, and Kaplan–Meier survival analysis

The gene expression profiles were obtained from the TCGA and GTEx databases. Differential expression was calculated with reads counts by DESeq2 after normalization. The cutoff was set as q < 0.05 and |log2FC| > 1. The protein expression profiles were obtained from the CPTAC databases. Z-values represent standard deviations from the median across samples for the given cancer type or stage. Log2 transformed spectral count ratio values were first normalized within each sample profile, then normalized across samples. Exon expression and isoform percentage were viewed and downloaded with UCSC Xena. PSI values of AS events in ovarian cancer were obtained from TCGASpliceseq (Ryan et al., 2016).

The analysis of the functional enrichment of differentially expressed genes was conducted using PANTHER, GO, and GSEA. GOplot, ggplot2 in R/Bioconductor 3.6.3, and GraphPad Prism 8 were used for plotting. An online Kaplan–Meier plotter database (http://kmplot.com/analysis/) was used to analyze the association between the mRNA expression levels of genes of interest and the survival information of patients with serous ovarian cancer (Jia et al., 2019). Cohorts of patients were split by median expression values through auto select cutoff. Survival analysis and Kaplan–Meier survival curves were performed in the R packages survival and survminer. High and low expression groups of our tissue microarray were defined by IHC staining score. All patients with overall survival or progression-free survival information were included.

### Statistical analysis

GraphPad Prism 8 and R (version 3.6.3) were used for statistical analysis. Student’s t-test and one-way analysis of variance were used to determine significant differences. Pearson’s correlation coefficient was used to determine the correlations between gene expression. The chi-square test was used to analyze the differences in clinical characteristics, and the log-rank test was used to detect differences in clinical prognosis. Cumulative distribution was analyzed by the Kolmogorov-Smirnov test. The results are presented as the means ± S.D. of three independent experiments. Statistical significance was considered as p < 0.05.

## AVAILABILITY

The RNA-seq, RIP-seq, and SpyCLIP data for this study are available for download from the Gene Expression Omnibus (GEO) repository (GSE183449, GSE183450, and GSE183451). The raw data for IP-MS are available in Table S3. The custom code used to analyze the data has been deposited at https://github.com/PrinceWang2018/BUD31_BCL2L12. The raw data and code are publicly available as of the date of publication. Any additional information required to reanalyze the data reported in this paper is available from the lead contact upon request.

## ACKNOWLEDGEMENTS

We thank the Translational Medicine Core Facility of Shandong University for consultation and instrument availability that supported this work.

